# Symbiont-mediated protection varies with wasp genotype in the *Drosophila melanogaster-Spiroplasma* interaction

**DOI:** 10.1101/691683

**Authors:** Jordan E Jones, Gregory D D Hurst

## Abstract

The ability of an insect to survive attack by natural enemies can be modulated by the presence of defensive symbionts. Study of aphid-symbiont-enemy interactions has indicated that protection may depend on the interplay of symbiont, host and attacking parasite genotypes. However, the importance of these interactions are poorly understood outside of this model system. Here, we study interactions within a *Drosophila* model system, in which *Spiroplasma* protect their host against parasitoid wasps and nematode attack. We examine whether the strength of protection conferred by *Spiroplasma* to its host, *Drosophila melanogaster* varies with strain of attacking *Leptopilina heterotoma* wasp. We perform this analysis in the presence and absence of ethanol, an environmental factor that also impacts the outcome of parasitism. We observed that *Spiroplasma* killed all strains of wasp. However, the protection produced by *Spiroplasma* following wasp attack depended on attacking wasp strain. A composite measure of protection, including both the chance of the fly surviving attack and the relative fecundity/fertility of the survivors, varied from a <4% positive effect of the symbiont following attack of the fly host by the Lh14 strain of wasp to 21% for the Lh-Fr strain in the absence of ethanol. Variation in protection provided was not associated with differences in the oviposition behaviour of the different wasp strains. We observed that environmental ethanol altered the pattern of protection against wasp strains, with *Spiroplasm*a being most protective against the Lh-Mad wasp strain in the presence of ethanol. These data indicate that the dynamics of the *Spiroplasma*-*Drosophila*-wasp tripartite interaction depend upon the genetic diversity within the attacking wasp population, and that prediction of symbiont dynamics in natural systems will thus require analysis across natural enemy genotypes and levels of environmental ethanol.

**Impact Summary:** Natural enemies – predators, parasites and pathogens – are a common source of mortality in animals, and this has driven the evolution of an array of mechanisms for preventing and surviving attack. Recently it has been observed that microbial symbionts form a component of insect defence against attack by pathogens and parasites. Whether an individual fly dies or lives following wasp attack, for instance, is partly determined by the presence or absence of *Spiroplasma* bacteria in the fly blood. The evolutionary biology of these ‘protective symbioses’ will in part depend on the specificity of defence – does *Spiroplasma* defend against all wasp strains equally, or does defence vary between wasp strains? We investigated this in the model insect, *Drosophila melanogaster*. We observed that the defensive symbiont killed all strains of wasps tested. However, the capacity of the symbiont to rescue the fly varied – *Spiroplasma* rescued the flies for some attacking wasp strains, but not for others. These data mean that the degree to which symbionts protect their host will depend on the wasp strains circulating in nature. Our results are important in terms of understanding the forces that promote symbiont mediated protection and understanding the origins of diversity of circulating wasp strains. Further, these data indicate enemy diversity and their interaction with protective symbionts should be included in evaluation of the efficiency of biocontrol programmes involving natural enemies.

## Introduction

All organisms face a threat from natural enemies and, in response, are typically able to defend themselves through a variety of protective mechanisms. In many species, the outcome of an encounter may be in part be determined by defensive symbionts within the host (and indeed offensive symbionts in the natural enemy) (Oliver *et al.* 2003; Scarborough *et al.* 2005; Xie *et al.* 2010, 2014; Jaenike *et al.* 2010; Lukasik *et al.* 2013; Dheilly *et al.* 2015; Mateos *et al.* 2016; Paredes *et al.* 2016; Ballinger & Perlman 2017). In insects, vertical transmission of bacterial symbionts places heritable symbionts into direct conflict with the natural enemies of their host. This conflict has driven the evolution of host protection in a number of symbiont clades, in a wide range of host species, against a diverse range of enemies. For example, microbial symbionts are known to provide protection against ssRNA viruses, nematodes, fungal pathogens and parasitic wasps (Oliver *et al.* 2003; Scarborough *et al.* 2005; Hedges *et al.* 2008; Teixeira *et al.* 2008; Xie *et al.* 2010, 2014; Jaenike *et al.* 2010; Lukasik *et al.* 2013; Mateos *et al.* 2016; Paredes *et al.* 2016; Ballinger & Perlman 2017).

Studies of defensive symbiosis are most well developed in aphid-symbiont-enemy interactions. For example, in the black bean aphid (*Aphis fabae*), the level of resistance conferred against the parasitoid (*Lysiphlebus fabarum*) is dependent on the interaction between the strain of defensive symbiont (*Hamiltonella defensa* carrying phage APSE) and the strain of the parasitoid, not the host itself (Oliver *et al.* 2009; Schmid *et al.* 2012; Cayetano & Vorburger 2013, 2015). Similarly, in the pea aphid (*Acyrthosiphon pisum*), protection against the entomopathogenic fungus, *Pandora neoaphidis*, is strongly dependent on the genotype-by-genotype interaction between the parasite and the defensive facultative symbiont, *Regiella insecticola* (Parker *et al.* 2017). Although these studies have demonstrated the importance of heritable microbes in mediating host-parasite specificity, the generality of these interaction terms is yet to be determined beyond the aphid system.

Regarded as a historically important model system for defence ecology and evolution, symbiont-mediated protection also occurs in the genus *Drosophila*. The facultative endosymbiont, *Spiroplasma*, can protect *Drosophila* against a range of endoparasitoid wasps. In *Drosophila hydei*, the native *Spiroplasma* strain Hy1 protects flies from the endoparasitoid wasp, *Leptopilina heterotoma* (Xie *et al.* 2010), although wasp attack survivors are found to have reduced fertility (Xie *et al*., 2011). Similarly, in *Drosophila melanogaster*, the *Spiroplasma* strain MSRO protects flies attacked by *Leptopilina boulardi* (Xie *et al.* 2014; Paredes *et al.* 2016; Ballinger & Perlman 2017), *Leptopilina victoriae* and *Ganapis xanthopoda* (Mateos *et al.* 2016). In *Drosophila neotestacea*, *Spiroplasma* confers tolerance against *Howardula* nematode worms, rescuing the fertility of female fly hosts (Jaenike *et al*., 2010).

Despite their importance as a model system, our understanding of *Spiroplasma*-mediated protection in *Drosophila* is limited in comparison to the equivalent aphid systems. Exploration of evolutionary dynamics is limited to the observation of the sweep of protective symbionts through North American *D. neotestacea* over time (Jaenike *et al*., 2010). More attention has been given to establishing the extent and molecular underpinnings of the defensive mechanisms. Variation in protective capacity against different parasitoid natural enemies has been observed. For example, whilst *Spiroplasma* strain MSRO is only very weakly able to rescue *D. melanogaster* flies parasitised by *L. heterotoma*, the same symbiont strain that increases fly survival by 50% against *L. boulardi* (Xie *et al.* 2014; Paredes *et al.* 2016; Ballinger & Perlman 2017). Defence is considered mechanistically to occur through a combination of RIP toxins secreted by the symbiont, and competition between symbiont and wasp for lipid (Paredes *et al*., 2016, Ballinger & Perlman 2017).

In this study, we determine whether the variation in *Spiroplasma*-mediated protection previously observed against different wasp species is reflected also in variation in protection against different strains of the same wasp species. This question is important as it determines whether the property of defence ascertained from a single wasp strain of a species describes symbiont-mediated defence generally against that species. Furthermore, we also determine whether the degree of protection and specificity to parasite strain are altered by the presence and absence of ethanol. Previous work has shown that environmental ethanol is an important determinant of the outcome of parasitoid wasp attack in *Drosophila*, with consumption of ethanol by infected larvae increasing mortality of wasp larvae growing within the hemocoel (Milan *et al.* 2012; Lynch *et al.* 2017). Determining variation in *Spiroplasma*-mediated protection across strains, and how this interacts with other protective mechanisms governed by the environment – such as ethanol-mediated protection, is key to understanding the dynamics of symbiont-mediated defence in natural populations.

To determine the extent to which wasp genotype affects the protective phenotype of *Spiroplasma* in *D. melanogaster*, a full factorial design was established including *Spiroplasma* infection status (positive and negative), ethanol (presence and absence) and wasp strain (one of three *L. heterotoma* strains). We combine survival data with data on the fertility of wasp attack survivors (compared to unattacked) to establish a protective index for each combination, which represents the first composite measure of symbiont-mediated protection obtained in any system to date. The size of flies that survived wasp attack and unattacked controls were also measured to determine stress experienced during larval development (stressed flies emerge at smaller size) (Miller & Thomas 2006). Finally, to determine whether any differences in protection conferred by *Spiroplasma* against wasps were due to variation in oviposition behaviour between the wasp strains, we also determined the number of wasp eggs/larvae per *Drosophila* larvae attacked by each wasp strain.

## Methods

### Insect strains and maintenance

*D. melanogaster* Canton-S flies with and without *Spiroplasma* MSRO-infected Red 42 were used. MSRO-infected Red 42 were originally collected in Brazil in 1997 and maintained in the lab in a Canton-S background in parallel to Canton-S control stock lacking *Spiroplasma*, from which males were derived each generation for MSRO line maintenance (Montenegro *et al.* 2000). These stocks both carried *Wolbachia* strain wmelCS, which occurs naturally and does not affect protection (Xie *et al.* 2014). It should be noted that all larvae from the *Spiroplasma* infected treatments are female due to the high efficiency of male-killing. However, there does not appear to be any differences in survival between the sexes against parasitoid wasp attack (Xie *et al.* 2014). All flies were maintained on Corn Meal Agar (10 g agarose, 85 g sugar, 60 g maize meal, 40 g autolysed yeast in a total volume of 1 L, to which 25 mL 10% Nipagin was added) at 25°C on a 12:12 light:dark cycle.

The *L. heterotoma* used were an inbred strain collected from Sainte Foy-lès-Lyon and la Voulte, France, a strain caught in Madeira, Portugal in March 2017, and the highly virulent inbred strain Lh14 used in previous studies, initially collected in Winters, California in 2002 (Schlenke *et al.* 2007). All wasp strains tested positive for *Wolbachia*. Wasp stocks were maintained on second instar Oregon-R larvae at 25°C on a 12:12 light:dark cycle. After emergence, wasps were maintained in grape agar vials supplemented with a flug moistened with honey water and allowed to mature and mate for 7 days prior to exposure to *D. melanogaster* L2 larvae.

### Preparing ethanol food

The wasp attack assay was performed in fly medium at 0% and 6% ethanol, which is within the normal range experienced by *D. melanogaster* larvae in nature (McKenzie & McKechnie 1979; Gibson *et al.* 1981). Medium was prepared by using the standard Corn Meal Agar recipe (above) with the exception of the quantity and concentration of Nipagin added (5 mL 50% w/v / 1 L of medium), to ensure the concentration of ethanol in the experimental vials was close to 0% and 6%. To prevent the evaporation of ethanol during the process, 200 mL of food was dispensed into 250 mL Duran bottles and allowed to cool to 45°C before 12 mL of 100% ethanol was added to the ethanol treatment bottles and homogenised. 6 mL of food was then dispensed into standard *Drosophila* vials and instantly covered with Parafilm to prevent ethanol evaporation before experimental larvae were transferred into the vials.

### Wasp attack assay

To ensure efficient vertical transmission of *Spiroplasma*, MSRO-infected Red 42 females were aged to at least ten days prior to egg laying. Flies were allowed to mate in cages and lay eggs on a grape Petri dish painted with live yeast for 24 h. Grape Petri dishes were incubated for a further 24 h to allow larvae to hatch. First instar larvae were picked from the grape plate into the experimental vials at 30 larvae per vial. Eight treatments were formed per wasp strain with approximately 10-15 replicate vials per treatment (1) Lh-S-EtOH-, (2) Lh-S-EtOH+, (3) Lh-S+ EtOH-, (4) Lh-S+ EtOH+, (5) Lh+ S-EtOH-, (6) Lh+ S-EtOH+, (7) Lh+ S+ EtOH-, (8) Lh+ S+ EtOH+. Five experienced female wasps and three male wasps were transferred into the wasp treatment vials. Flugs were used to bung vials to reduce ethanol evaporation. Adult wasps were allowed to parasitise for 2 days before being removed. All vials were maintained at 25°C on a 12:12 light:dark cycle. For each vial, the number of pupae, emerging flies and emerging wasps were recorded.

### Measuring fertility

To determine the degree to which survivors of wasp attack were impacted by wasp attack, the average daily emerged offspring of *Spiroplasma* infected survivors (“Exposed”) and *Spiroplasma* infected flies which did not undergo wasp attack (“Unexposed”) was measured from both the 0% and 6% ethanol treatments. Only fly survivors which underwent attack from the *L. heterotoma* Lh-Fr and Lh-Mad strain were used as there were few survivors from attack of the Lh14 strain of *L. heterotoma* and very low numbers also from the *Spiroplasma* uninfected wasp attacked group.

To this end, adult female flies from the wasp attack assay were retained on eclosion, and stored in vials containing sugar yeast medium (20 g agarose, 100 g sugar, 100 g autolysed yeast in a total volume of 1 L, to which 30 mL 10% Nipagin w/v propionic acid was added) at mixed ages. A week after emergence commenced, approximately 45 female flies from each of the *Spiroplasma* treatments were placed individually into an ASG vial with two Canton-S males with a single yeast ball and allowed to mate. These flies were transferred onto fresh ASG vials each day for five days. Female fertility was measured as the average number of daughters produced over four days (day 2-5), with F1 flies given two weeks to emerge to ensure every fly had emerged before counting. Females which did not produce any daughters were considered infertile.

### Measuring wing size

Body size as adult measures the stress experienced by flies during development, with many stresses (density, ethanol) resulting in smaller adult flies (Miller & Thomas 2006; Castañeda & Nespolo 2013). To determine whether wasp attack affected female body size, wing size was used as a proxy, as these factors are known to be highly correlated in *Drosophila* (Robertson & Reeve 1952). To this end, the left wings of individual flies from the experiment above were removed using forceps under a microscope (right wings were used if left wings were damaged) and mounted flat onto a glass microscope slide. A photograph was taken of each wing using a microscope mounted camera using GXCapture-O software (6.9v). Using ImageJ software (1.49v, US National Institutes of Health, USA), the area of the wing was determined by locating the coordinates of the six wing landmarks as defined in Gilchrist and Partridge (2001) and calculating the interior area of the polygon created. A scale slide was used to transform all wing measurements into millimetre square units. All photos where the landmarks were not clearly visible were not measured and excluded from the analysis.

### Wasp strain oviposition

To determine whether the differences in fly survival were due to differences in wasp oviposition behaviour, we compared the number of wasp eggs and larvae per fly larva among the three wasp strains (Lh-Fr, Lh14 and Lh-Mad). In addition, we determined whether wasp oviposition differed between *Spiroplasma* positive and negative fly larvae. To this end, we followed the same protocol as the wasp assay, except the no-wasp control and 6% ethanol treatment was omitted. Immediately after wasp removal, approximately 5 fly larvae from each of the five replicate vials were dissected under a microscope to count the number of wasp eggs and/or larvae present.

### Statistical analysis

All statistical analyses were performed using the statistical software R, version 3.5.0 (R Core Team, 2018). All data was tested for normality.

#### Fly survival

A generalized linear model with binomial errors was used to test the effect of symbiont, ethanol and wasp strain (where appropriate) on fly larva-to-adult survival. First, the maximal model was tested including all factors and their interactions. The model was simplified stepwise until only significant factors remained.

#### Wasp survival

Due to extreme separation in wasp survival between symbiont treatments (*Spiroplasma* positive treatments had 0 wasp survival), a Bayesian generalized linear model (‘bayesglm’ function in the ‘arm’ package; Gelman *et al*., 2018) with binomial errors was used to test the effect of fly symbiont status, ethanol and wasp strain on wasp larva-to-adult survival. The model was simplified stepwise until only significant factors remained.

#### Fertility

A generalized linear model with binomial errors was used to test the effect of wasp attack and ethanol on the proportion of flies fertile. Linear models were used to test the effect of wasp attack and ethanol on the average number of daughters produced from fertile flies. First the maximal model was tested including all factors and their interaction. The model was simplified stepwise until only significant factors remained.

#### Composite measure of protection

We calculated a Protective Index (PI), comparing the survival and fecundity of *Spiroplasma* infected flies in the presence/absence of a given strain of wasp. PI was calculated as the ratio of p(survival) × p(fertile) × fecundity of fertile individuals for attacked vs unattacked *Spiroplasma* infected flies and reflects the benefit of *Spiroplasma* in the face of wasp attack. Credible intervals for PI were calculated through simulation. By assuming prior probability distributions for each parameter (Survival probability = beta distribution; Fertility probability = beta distribution; Fecundity = normal distribution), the ‘rbeta’ and ‘rnorm’ functions were used to calculate 95% credible intervals for PI. The simulation data was also used to establish the posterior probability of PI differing between attacking wasp strains.

#### Wing size

Wing area data was transformed using Box-Cox transformations (Crawley 2007). Linear models were used to test the effect of wasp attack and ethanol on wing size. First the maximal model was tested including all factors and their interaction. The model was simplified stepwise until only significant factors remained.

#### Differential oviposition

A generalized linear model with a Poisson error structure was used to test the effect of wasp strain and fly *Spiroplasma* infection status on the number of wasp eggs/larvae found laid into fly larva. First the maximal model was tested including all factors and their interaction. The model was simplified stepwise until only significant factors remained.

## Results

### Fly survival and wasp success

In the absence of *L. heterotoma*, mean larva-to-adult fly survival was >69% across all treatments (Fig.1). There was no significant effect of *Spiroplasma* or ethanol, nor a significant interaction between *Spiroplasma* and ethanol on fly larva-adult survival.

**Fig. 1:**
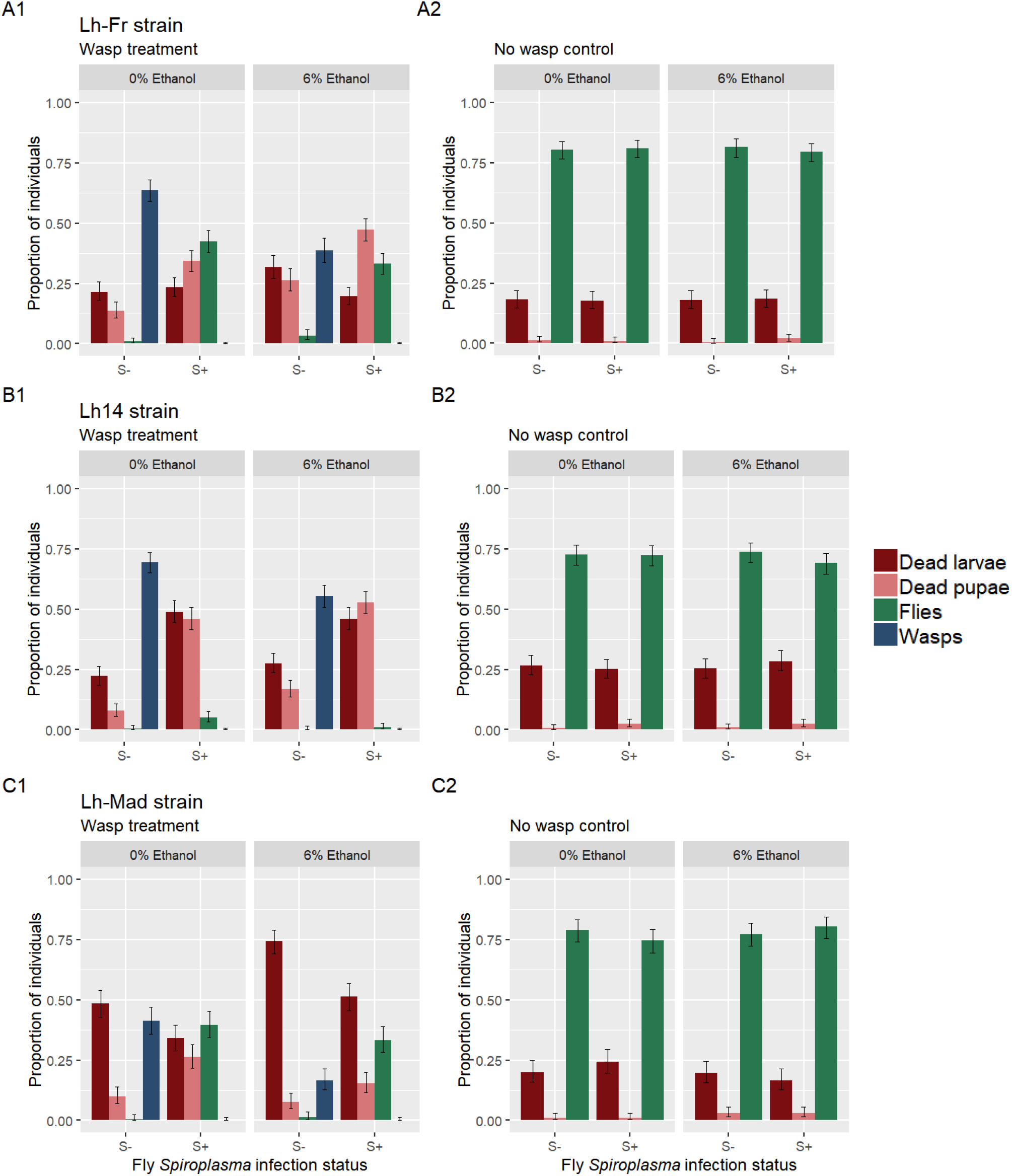
Proportion of dead larvae (red), dead pupae (pink), emerging flies (green) and emerging wasps (blue) for *Spiroplasma*-infected and uninfected *Drosophila melanogaster* attacked by three different *Leptopilina heterotoma* strains in 0% and 6% environmental ethanol. Error bars represent 95% binomial confidence intervals. (Lh-Fr: n= 15 replicate vials per treatment except S-EtOH+ Lh+ (12 vials) and S-EtOH+ Lh-(13 vials), Lh-Mad: n = 10 replicate vials per treatment, Lh14: n = 15 replicate vials per treatment).

In the presence of *L. heterotoma*, fly *Spiroplasma* infection had a significantly strong and positive effect on fly larva-to-adult survival (*P* < 0.001; Fig. 1). The effect of *Spiroplasma* on fly larva-to-adult survival depended on the strain of attacking parasitoid, which was reflected in a significant interaction between *Spiroplasma* and wasp strain (*P* < 0.05). *Spiroplasma* provided almost no protection against the Lh14 strain of *L. heterotoma*, increasing fly larva-to-adult survival slightly from <1% to 5.11%. *Spiroplasma* did however, provide strong protection against the Lh-Fr and Lh-Mad wasp strains, increasing fly larva-to-adult survival from <1% to 42.4% and 39.7% respectively. Wasp strain itself had a significant effect on fly larva-adult survival (*P* < 0.001).

The presence of ethanol had a weak, albeit significant positive effect on fly larva-adult survival in the presence of wasps (*P* < 0.01; Fig. 1). However, the effect of ethanol differed between the strains of attacking *L. heterotoma*, which was reflected in a significant interaction between ethanol and wasp strain (*P* < 0.05). Specifically, the presence of ethanol in the absence of *Spiroplasma* reduces fly larva-to-adult survival against the Lh14 *L. heterotoma* strain from 0.45% to 0.22%, yet slightly increases fly larva-to-adult survival against the Lh-Fr strain from 0.89% to 3.33% and the Lh-Mad strain from 0.33% to 1.33%. There was also a significant interaction between *Spiroplasma* and ethanol (*P* < 0.001; Fig. 1), with the presence of ethanol reducing the effect of *Spiroplasma*-mediated fly larva-to-adult survival across all three wasp strains (% decrease; Lh-Fr = 22%, Lh14 = 78%, Lh-Mad = 16%). The interaction between *Spiroplasma*, wasp strain and ethanol was not found to be significant.

Wasp success was strongly negatively affected by fly *Spiroplasma* infection, with the presence of *Spiroplasma* completely preventing the emergence of wasps across all *L. heterotoma* strains in both the presence and absence of ethanol. In the absence of *Spiroplasma*, the presence of ethanol had a significantly negative effect on wasp success (*P* < 0.001; Fig. 1). However, the effect of ethanol depended on the strain of attacking *L. heterotoma*, reflected in a significant interaction between ethanol and wasp strain (*P* = 0.014). Ethanol reduced wasp success by 40%, 21%, and 60% across the Lh-Fr, Lh14 and Lh-Mad strains respectively. Wasp success was also significantly affected by the strain of wasp (*P* < 0.001). There was no significant interaction between ethanol and *Spiroplasma*, *Spiroplasma* and wasp strain, nor a significant 3-way interaction on wasp success.

### Female fertility

#### Proportion fertile

For both Lh-Mad and Lh-Fr attacking wasp strains, *Spiroplasma*-infected individuals that survived wasp attack were observed to have reduced fertility, measured as the proportion of females able to produce progeny (Fig. 2).

**Fig. 2:**
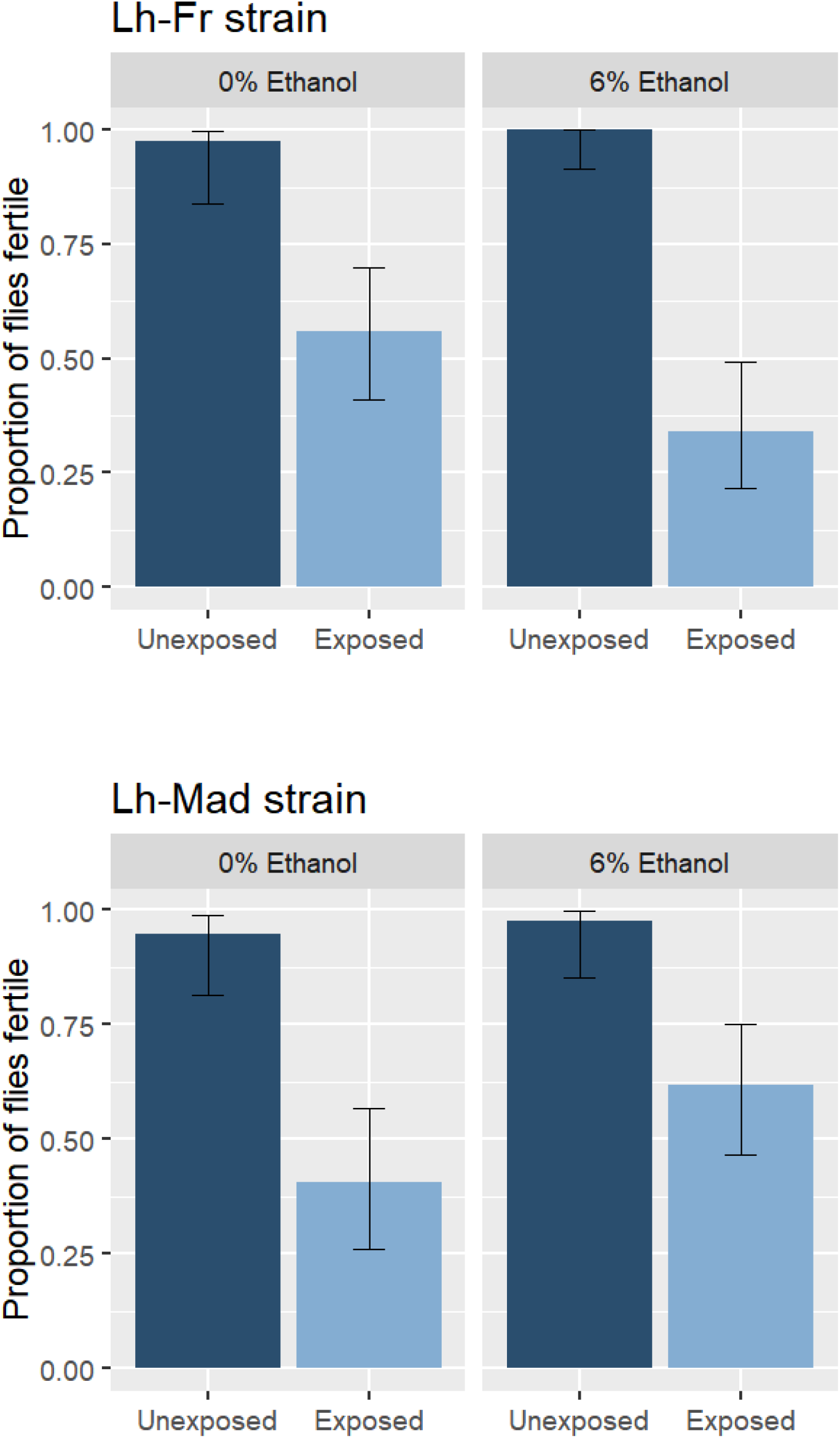
The proportion of *Spiroplasma*-infected *Drosophila melanogaster* females considered fertile after exposure to *Leptopilina heterotoma* (Lh-Fr and Lh-Mad strain) and unexposed controls developed through 0% and 6% ethanol medium. Dark blue bars indicate unexposed controls and light blue bars represent wasp exposed. Error bars represent 95% binomial confidence intervals. Lh-Fr strain: 87 wasp-attacked (0% EtOH n=43 and 6% EtOH n=44) and 80 non-wasp attacked (0% EtOH n=39 and 6% EtOH n=41); Lh-Mad strain: 79 wasp-attacked (0% EtOH n=37 and 6% EtOH n=42) and 81 non-wasp attacked (0% EtOH n=38 and 6% EtOH n=43).

For attack with the Lh-Fr strain of wasp, there was a significant effect of wasp attack on the proportion of flies which were found to be fertile (*P* < 0.001; Fig. 2). The proportion of *D. melanogaster* considered fertile following wasp-attack was reduced by 55% compared to control non-attacked *D. melanogaster*. There was no significant effect of ethanol (*P* = 0.167), nor a significant interaction between ethanol and wasp attack (*P* = 0.117).

For attack with the Lh-Mad strain, there was a significant effect of wasp attack on the proportion of flies found to be fertile (*P* < 0.001; Fig. 2). The proportion of *D. melanogaster* considered fertile following wasp-attack was reduced by 46% compared to control non-attacked *D. melanogaster*. There was no significant effect of ethanol (*P* = 0.083), nor a significant interaction between ethanol and wasp attack (*P* = 0.987).

#### Number of daughters produced

In both cases, *Spiroplasma*-infected individuals that survived wasp attack and were fertile were observed to produce fewer daughters compared to fertile, unattacked controls (Fig. 3).

**Fig. 3:**
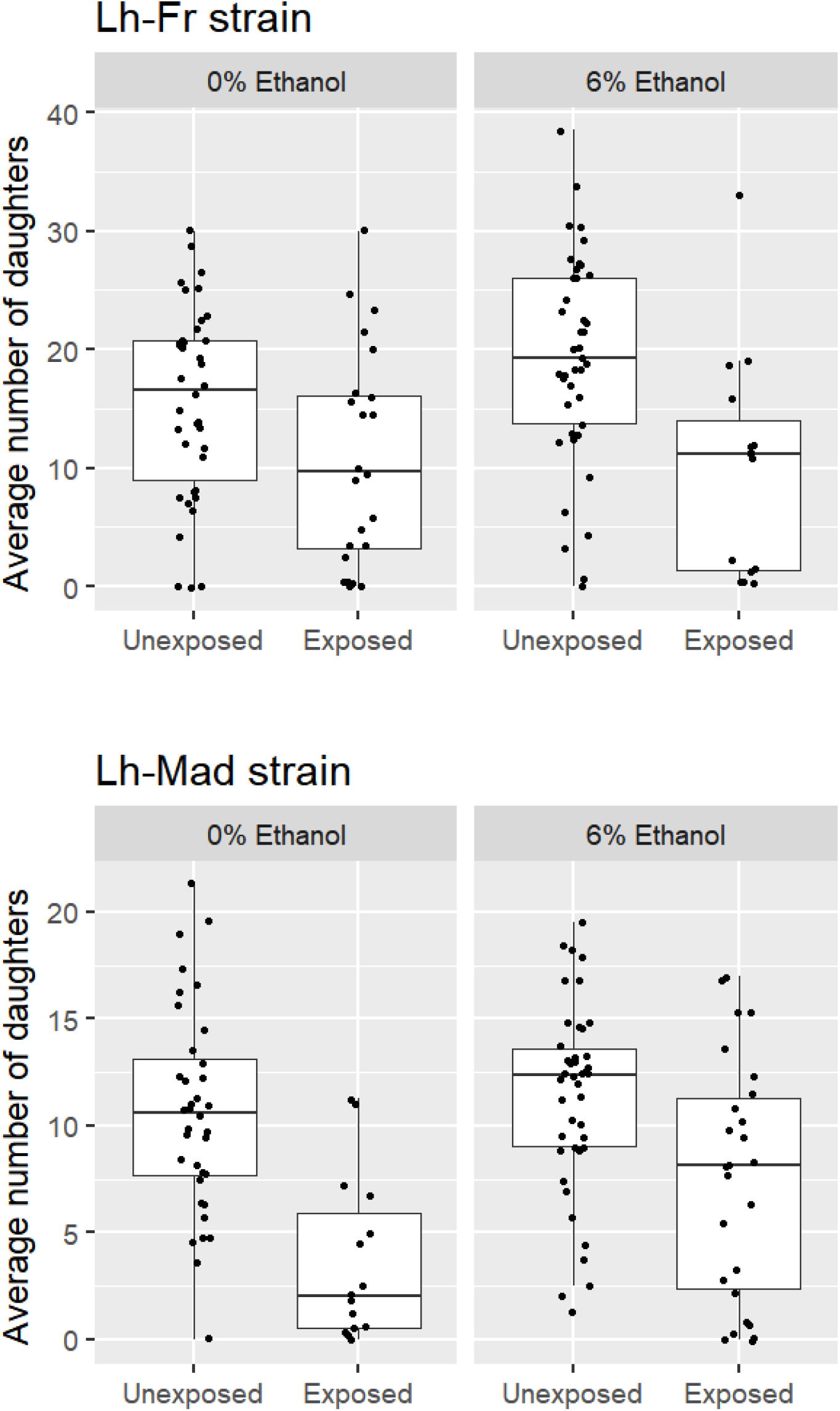
The average number of daughters produced by fertile *Spiroplasma*-infected female *Drosophila melanogaster* exposed to *Leptopilina heterotoma* (Lh-Fr and Lh-Mad strain) and unexposed controls developed through 0% and 6% ethanol medium. The box plots display the upper and lower quartiles, the median and the range. Points represent each measurement obtained. Lh-Fr strain: 39 wasp-attacked (0% EtOH n=24 and 6% EtOH n=15) and 79 non-wasp attacked (0% EtOH n=38 and 6% EtOH n=41); Lh-Mad strain: 41 wasp-attacked (0% EtOH n=15 and 6% EtOH n=26) and 78 non-wasp attacked (0% EtOH n=36 and 6% EtOH n=42).

For attack with the Lh-Fr strain, wasp attack significantly reduced the average number of daughters produced with protected wasp attacked *D. melanogaster* averaging ~39% fewer than control unattacked *D. melanogaster* (Mean ± SE = 10.6 ± 1.44 daughters for attacked flies vs. 17.5 ± 0.969 daughters for control flies; *P* < 0.001; Fig. 3). There was no significant effect of ethanol (*P =* 0.067), nor a significant interaction between ethanol and wasp attack (*P =* 0.190).

For attack with the Lh-Mad strain, wasp attack also significantly reduced the average number of daughters produced with wasp attacked protected *D. melanogaster* averaging ~45% fewer than control *D. melanogaster* (Mean ± SE = 6.11 ± 0.832 daughters for attacked flies vs. 11.0 ± 0.521 daughters for control flies; *P* < 0.001; Fig. 3). There was no significant effect of ethanol (*P =* 0.143), nor a significant interaction between ethanol and wasp attack (*P =* 0.097).

#### Overall protection

Taking into account the survival, proportion of adults fertile, and the fecundity of wasp attack survivors, compared to unexposed *Spiroplasma* infected controls, a protection index (PI) was calculated as the product of fly survival x p(fertile) x fecundity of exposed vs unexposed *Spiroplasma* infected flies (this metric assumes complete mortality from wasps in the absence of *Spiroplasma*, which is approximately true as <1% of individuals tested survived wasp attack). In the absence of ethanol, this estimated a protection index of 21%, and 8% against the Lh-Fr and Lh-Mad strains respectively (Table 1). The posterior probability that the protection index for *Spiroplasma* against the Lh-Fr strain is greater than the protection index against the Lh-Mad strain is 0.99. In contrast, the PI in the presence of ethanol were 7% and 17% against Lh-Fr and Lh-Mad wasp strains respectively (Table 1). The posterior probability that the protection index for *Spiroplasma* against the Lh-Mad strain is greater than the protection index against the Lh-Fr strain in the presence of ethanol is 0.99. With no fecundity measure available for Lh14 (due to insufficient survivors), we assume the estimate of protection to be less than the survival value.

**Table 1:** The overall protection conferred by *Spiroplasma* against the Lh-Fr, Lh14 and Lh-Mad *Leptopilina heterotoma* strains in *Drosophila melanogaster* in the presence and absence of ethanol. Exposed S− = wasp attacked *Spiroplasma* uninfected flies; Exposed S+ wasp attacked *Spiroplasma* infected flies; Unexposed S+ *Spiroplasma* infected flies not attacked. Protective Index is calculated as [p(survival) x p(fertile) x fecundity of fertile individuals] of exposed vs unexposed individuals with credible intervals calculated as given in methods.

| Wasp strain | Treatment | Fly Survival<br>(binomial 95% CI intervals<br>(lower, upper)) | Proportion fertile<br>(binomial 95% CI intervals<br>(lower, upper)) | Fecundity<br>measure<br>± SE | Estimated<br>protective index<br>(95% Credible<br>interval (lower,<br>upper)) |
| --- | --- | --- | --- | --- | --- |
| **Lh-Fr** | Exposed S- | <0.01 (0.0033 - 0.023) | N/A | N/A |  |
|  | Exposed S+ | 0.42 (0.38 - 0.47) | 0.56 (0.41 - 0.70) | 10.9 ± 1.83 |  |
|  | Unexposed control S+ | 0.81 (0.77 - 0.84) | 0.97 (0.84 - 0.99) | 15.6 ± 1.31 | 0.21 (0.12, 0.33) |
| **Lh14** | Exposed S- | <0.01 | N/A | N/A |  |
|  | Exposed S+ | 0.05 | N/A | N/A |  |
|  | Unexposed control S+ | 0.72 | N/A | N/A | <0.036 |
| **Lh-Mad** | Exposed S- | <0.01 (0.00047 - 0.023) | N/A | N/A |  |
|  | Exposed S+ | 0.40 (0.34 - 0.45) | 0.41 (0.26 - 0.57) | 3.65 ± 0.992 |  |
|  | Unexposed control S+ | 0.75 (0.69 - 0.79) | 0.95 (0.81 - 0.99) | 10.6 ± 0.801 | 0.08 (0.032, 0.15) |

b) In the presence of 6% ethanol
| **Wasp strain** | **Treatment** | **Fly Survival**<br>**(binomial 95% CI intervals**<br>**(lower, upper))** | **Proportion fertile**<br>**(binomial 95% CI intervals**<br>**(lower, upper))** | **Fecundity**<br>**measure**<br>**± SE** | **Estimated**<br>**protective index**<br>**(95% Credible**<br>**intervals (lower,**<br>**upper))** |
| --- | --- | --- | --- | --- | --- |
| **Lh-Fr** | Exposed S- | 0.03 (0.019, 0.058) | N/A | N/A |  |
|  | Exposed S+ | 0.33 (0.29, 0.38) | 0.34 (0.22, 0.49) | 9.98 ± 2.41 |  |
|  | Unexposed control S+ | 0.80 (0.76, 0.83) | 1 (0.91, 1.00) | 19.2 ± 1.38 | 0.07 (0.033, 0.13) |
| **Lh14** | Exposed S- | <0.01 | N/A | N/A |  |
|  | Exposed S+ | 0.01 | N/A | N/A |  |
|  | Unexposed control S+ | 0.69 | N/A | N/A | <0.007 |
| **Lh-Mad** | Exposed S- | 0.01 (0.0050, 0.035) | N/A | N/A |  |
|  | Exposed S+ | 0.33 (0.28, 0.39) | 0.62 (0.47, 0.75) | 7.53 ± 1.10 |  |
|  | Unexposed control S+ | 0.80 (0.75, 0.84) | 0.98 (0.85, 1.00) | 11.3 ± 0.687 | 0.17 (0.11, 0.26) |

#### Wing size

In both cases, *Spiroplasma*-infected individuals that survived wasp attack, were smaller compared to unattacked *Spiroplasma* infected individuals (Fig. 4).

**Fig. 4:**
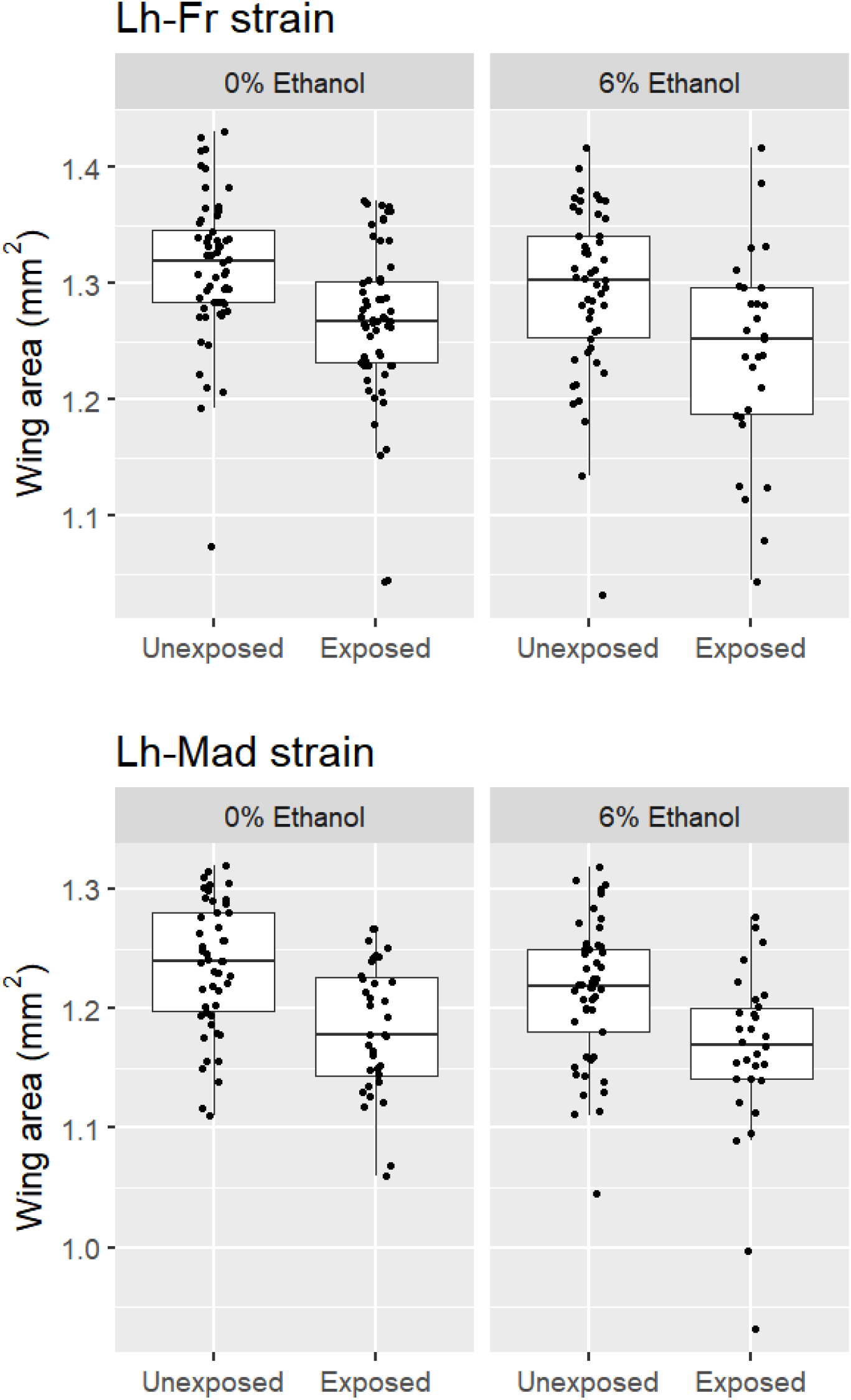
The wing area (mm^2^) of *Spiroplasma*-infected female *Drosophila melanogaster* exposed to *Leptopilina heterotoma* (Lh-Fr and Lh-Mad strain) and unexposed controls developed through 0% and 6% ethanol medium. The box plots display the upper and lower quartiles, the median and the range. Points represent each measurement obtained. Lh-Fr: 89 wasp-attacked (0% EtOH n=60 and 6% EtOH n=29) and 110 non-wasp attacked (0% EtOH n=60 and 6% EtOH n=50); Lh-Mad strain: 66 wasp-attacked (0% EtOH n=36 and 6% EtOH n=30) and 99 non-wasp attacked (0% EtOH n=50 and 6% EtOH n=49).

For attack with the Lh-Fr wasp strain, wasp attack strongly reduced wing size, with the wings of wasp attacked female *D. melanogaster* on average 0.04 mm^2^ (3%) smaller than unattacked *D. melanogaster* (Mean ± SE = 1.26 ± 0.008 mm^2^ for attacked flies vs. 1.30 ± 0.006 mm^2^ for unattacked flies; *P* < 0.001; Fig. 4). Ethanol reduced wing size, with the wing size of *D. melanogaster* reared in ethanol on average 0.02 mm^2^ (1.5%) smaller than *D. melanogaster* reared in the absence of ethanol (Mean ± SE = 1.27 ± 0.008 mm^2^ for flies reared in 6% ethanol vs. 1.29 ± 0.007 mm^2^ for control flies; *P* = 0.038; Fig. 4). There was no significant interaction between ethanol and wasp attack on wing size (*P* = 0.980).

For attack with the Lh-Mad wasp strain, wasp attack also had a highly significant effect on wing size, with the wing size of wasp attacked female *D. melanogaster* on average 0.05 mm^2^ (4%) smaller than control *D. melanogaster* (Mean ± SE = 1.17 ± 0.006 mm^2^ for attacked flies vs. 1.22 ± 0.006 mm^2^ for unattacked flies; *P* < 0.001; Fig. 4). There was no effect of ethanol (*P* = 0.053), nor a significant interaction between ethanol and wasp attack on wing size (*P* = 0.634).

#### Wasp oviposition

The average number of wasp eggs laid into a fly larva across a 48 h period of parasitisation was >1 but <2 for all treatments (Fig. 5). There was no significant effect of wasp strain (*P* = 0.087) or fly *Spiroplasma* infection status (*P* = 0.217), nor a significant interaction between wasp strain and fly *Spiroplasma* infection (*P* = 0.716) on the number of wasp eggs laid into fly larva.

**Fig. 5:**
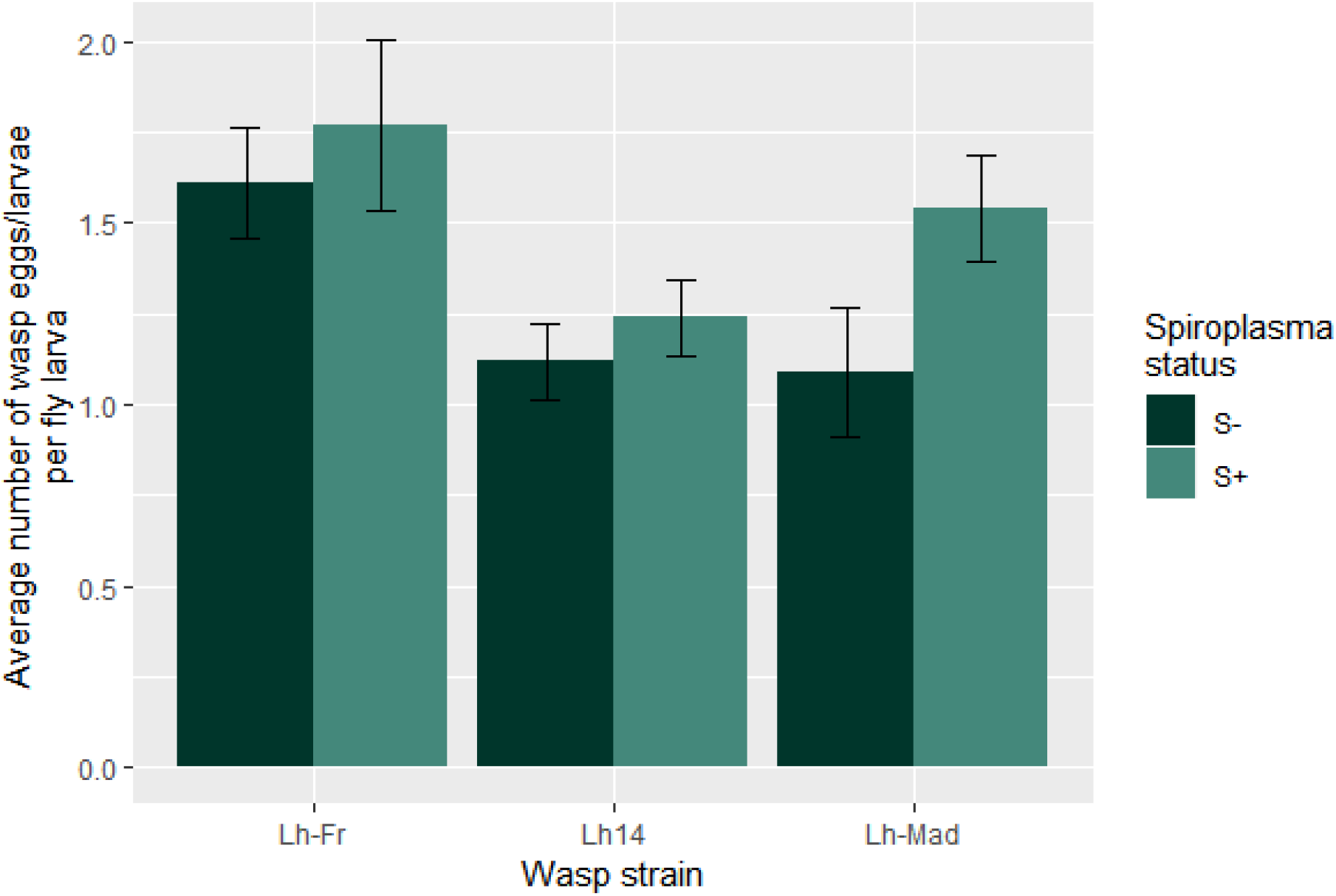
The average number of wasp eggs/larvae in *Spiroplasma* positive and negative *Drosophila melanogaster* larvae following 48 h of parasitisation by three strains of *Leptopilina heterotoma* (Lh-Fr: S-n= 23, S+ n=26; Lh14: S-n= 25, S+ n=25; Lh-Mad: S-n= 23, S+ n=24). Dark green bars indicate *Spiroplasma* negative individuals and light green bars represent *Spiroplasma* positive individuals. Error bars depict ± SE.

## Discussion

It is now recognised that the outcome of natural enemy attack can be determined by the presence or absence of defensive heritable symbionts. Beyond their presence, the outcome of these interactions can also depend on the genotypes of all players: symbiont, host and enemy. However, the specificity of symbiont-mediated defence has only been explored within the aphid system. Previous work has found wasp species to be an important component of *Spiroplasma*-mediated protection in *Drosophila*, with *Spiroplasma* able to protect against some wasp species, but not others (Mateos *et al.* 2016). Protection against *L. heterotoma*, using strain Lh14, for instance, is considered weak or absent in three previous studies (Xie *et al.* 2014; Paredes *et al.* 2016; Ballinger & Perlman 2017). In this study, we examined whether protection against *L. heterotoma* wasps varied with wasp strain. Protection against the Lh14 wasp strains was observed at the low level previously recorded. In contrast, substantial protection was exhibited against the other strains of *L. heterotoma*. The overall protection gained by harbouring *Spiroplasma* against the Lh-Fr, Lh-Mad and Lh14 *L. heterotoma* was approximately 21%, 8% and <4% respectively, measured in the absence of environmental ethanol. Thus, *Spiroplasma* is protective against *L. heterotoma*, but the degree of protection is wasp strain dependent.

The differences in protective index afforded by *Spiroplasma* against different wasps strains arose through both effects on survival in response to wasp attack (Lh14 attack kills flies notwithstanding *Spiroplasma* presence, whereas *Spiroplasma* rescues flies attacked by Lh-Mad/Lh-Fr strains) and through differences in fertility/fecundity (between flies surviving attack by Lh-Fr and Lh-Mad strains). Thus, we conclude the protection afforded by *Spiroplasma* against *L. heterotoma* is dependent on *L. heterotoma* genotype, and that the differences observed are a product of both fly survival and survivor fertility differences. We would note that whilst impacts on the fertility/fecundity of ‘protected’ survivors of attack is noted in some cases of defensive symbiosis (Xie *et al.* 2011; Vorburger *et al.* 2013), these metrics have not previously been included in models of relative protection against different enemy strains/species. Our data indicate that a complete model of protection dynamics may require measurement and inclusion of these parameters.

The mechanistic processes that determine the degree to which *Spiroplasma* affords protection against different wasp genotypes are uncertain. Whilst the presence of *Spiroplasma* completely prevented any wasps from emerging in all cases, the degree to which this rescued their fly host varied. Flies could be seen developing in the pupal cases in the majority of cases, but it was variation in eclosion to adult that underlies differential fly survival in response to the different genotypes of wasp. From the wasp differential oviposition assay, we can reject the hypothesis that the observed differences are due to differential oviposition behaviour across wasp strains.

One possible cause of variation in outcome is differential susceptibility of the wasp strains to *Spiroplasma*-produced RIP toxins. *Spiroplasma* produce RIP toxins which act against wasp ribosomes, causing wasp developmental failure (Ballinger & Perlman 2017). The differences in protection could be due to differences of the effect of these toxins on the wasp (Garcia-Arraez *et al.* 2019). Another explanation for the difference in survival could be variation in wasp virulence. It is possible that *Spiroplasma* infection is insufficient to rescue flies against more virulent wasp strains which better evade other endogenous defences within the fly. This effect could be driven by intraspecific venom variation amongst wasps (Colinet *et al.* 2013), which are transferred along with eggs to supress the host immune system to bypass nuclear encoded defences.

The protection offered by *Spiroplasma* against wasp strains is modified by the presence of environmental ethanol exposure during the larval phase. In contrast to assays where ethanol was absent, protection in the presence of ethanol is strongest against the Lh-Mad strain of wasp, and less strong against the Lh-Fr strain, with protection absent against Lh14. Against the Lh-Fr wasp strain, ethanol had a negative effect on the overall *Spiroplasma*-mediated fly protection, reducing protection from 21% to 7%. In contrast, ethanol had a positive effect on the overall protection against the Lh-Mad wasp strain, increasing overall protection from 8% to 17%, mainly due to the presence of ethanol reducing the negative effect of wasp attack on survivor fertility. In all cases, ethanol was detrimental to fly survival upon wasp attack. These results indicate that the interaction between *Spiroplasma*-mediated protection and ethanol protection is dependent on the genotype of the attacking wasp.

The data presented here have significant implications for the evolutionary and ecological dynamics of the *Spiroplasma*-*Drosophila*-wasp tripartite interaction in natural populations. From the perspective of the symbiont, the fitness benefit of protection is dependent upon wasp genotype, and thus the degree to which wasp attack drives the symbiont to higher prevalence will depend on the profile of the wasp population. In contrast, the observation that wasp emergence is zero in the presence of the symbiont in all three cases implies that the symbiont will not select upon the wasp population directly, although it will decrease the size of this population.

Environmental ethanol, which modulates wasp attack outcome, is likely to be less important than *Spiroplasma*-mediated protection in terms of determining wasp success. In contrast to other lab studies (Milan *et al.* 2012; Kacsoh *et al.* 2013; Lynch *et al.* 2017), here we estimated only a small magnitude of protection afforded by ethanol alone. Possible reasons for the disparity include variation in fly strains (Canton-S here, Oregon-R in other studies) and differences in experimental protocols (e.g, the period of exposure). Nevertheless, ethanol did determine the relative protective benefit of *Spiroplasma* against different wasp strains. Thus, the presence/absence of ethanol melds with the genetic makeup of the wasp population to determine protection accorded by *Spiroplasma*, and ultimately therefore is predicted to impact *Spiroplasma* dynamics.

In summary, our work has extended the aphid synthesis to *Drosophila*, and indicates symbiont mediated protection appears generally to depend on the genotype of the attacking wasp species. Further the environment (in this case ethanol) may modulate protection. More widely, it will be important not to disregard other protective mechanisms and their interaction when predicting the ecological dynamics of symbiont-mediated protection in this model system. Indeed, how *Spiroplasma*-mediated protection is predicted to interact with *Drosophila’s* own innate immunity (and more widely, host genetic background) requires further investigation. Beyond this, parallels with studies of aphids indicate that symbiont genotype and environment should be considered. Thermal environment, for instance, commonly affects symbiotic phenotype, and low temperatures are known ablate *Spiroplasma* male-killing (Anbutsu *et al.* 2008). Thus, whilst our study indicates the presence of complex interaction terms in this tripartite interaction, the full extent of these awaits resolution.

## Acknowledgements

We would like to thank Dr Todd Schlenke for providing the Lh14 wasp strain and Dr Fabrice Vavre for providing the Lh-Fr strain. We would also like to thank Dr Steve Parratt and Ben Walsh for collecting the wasp strain from Madeira (Lh-Mad). We thank Dr Stephen Cornell and Prof Andy Fenton for their help on the analysis and, Dr Ewa Chrostek and Dr Andrea Betancourt for their comments on the manuscript. This project was supported by funding from the NERC (Studentship to JJ, grant number NE/L002450/1).

## Author contributions

JJ and GH conceived and designed the study, JJ collected and analysed the data, JJ and GH drafted the initial version of the manuscript and contributed to later versions of the manuscript.

## Conflict of interest

The authors declare no conflicts of interest.

## Data accessibility

Data generated and analysed during this study are available at figshare (https://doi.org/10.6084/m9.figshare.c.4551422.v1).

## Notes

https://doi.org/10.6084/m9.figshare.c.4551422.v1

